# MRIO: The Magnetic Resonance Imaging Acquisition and Analysis Ontology

**DOI:** 10.1101/2023.08.04.552020

**Authors:** Alexander Bartnik, Lucas M. Serra, Mackenzie Smith, William D. Duncan, Lauren Wishnie, Alan Ruttenberg, Michael G. Dwyer, Alexander D. Diehl

**Affiliations:** Buffalo Neuroimaging Analysis Center, Department of Neurology, Jacobs School of Medicine and Biomedical Sciences, University at Buffalo, Buffalo, NY, USA; Department of Biomedical Informatics, Jacobs School of Medicine and Biomedical Sciences, University at Buffalo, Buffalo, NY, USA; University of Florida, College of Dentistry, Gainesville, FL, USA

**Keywords:** biological ontologies, magnetic resonance imaging, neuroinformatics, neuroimaging, Translational Research, Biomedical

## Abstract

**Objective:** Magnetic resonance imaging of the brain is a useful tool in both the clinic and research settings, aiding in the diagnosis and treatments of neurological disease and expanding our knowledge of the brain. However, there are many challenges inherent in managing and analyzing MRI data, due in large part to the heterogeneity of data acquisition.

**Materials and Methods:** To address this, we have developed MRIO, the Magnetic Resonance Imaging Acquisition and Analysis Ontology.

**Results:** MRIO provides well-reasoned classes and logical axioms for the acquisition of several MRI acquisition types and well-known, peer-reviewed analysis software, facilitating the use of MRI data. These classes provide a common language for the neuroimaging research process and help standardize the organization and analysis of MRI data for reproducible datasets. We also provide queries for automated assignment of analyses for given MRI types.

**Discussion:** MRIO aids researchers in managing neuroimaging studies by helping organize and annotate MRI data and integrating with existing standards such as Digital Imaging and Communications in Medicine and the Brain Imaging Data Structure, enhancing reproducibility and interoperability. MRIO was constructed according to Open Biomedical Ontologies Foundry principals and has contributed several terms to the Ontology for Biomedical Investigations to help bridge neuroimaging data to other domains.

**Conclusion:** MRIO addresses the need for a “common language” for MRI that can help manage the neuroimaging research, by enabling researchers to identify appropriate analyses for sets of scans and facilitating data organization and reporting.

## Introduction

Magnetic resonance imaging (MRI) is a biomedical imaging technique used to noninvasively visualize the internal structure of tissue in three-dimensional space. Because of this, MRI is well suited for studying the human brain *in vivo*, and has been employed in both clinical and research settings. Clinically, MRI is used to aid diagnosis of neurological diseases by helping identify the location and extent of pathology. Additionally, researchers can use MRI biomarkers to develop personalized treatments for neurological diseases^1^ and gain a deeper understanding of the brain’s structure, function, and connectivity.^2,3^ As the use of MRI increases, so does the amount of neuroimaging data produced.

While leveraging these data should lead to new discoveries and improve clinical outcomes, organizing and annotating it requires significant effort. Hence, there is a need for a “common language” governing the acquisition and analysis of MRI data. For many years, biomedical imaging has used the Digital Imaging and Communications in Medicine (DICOM) standard^4^ for providing a system of reporting the acquisition, storage, and transmission of data. DICOM is recognized by the International Organization for Standardization (ISO) as the official standard used for biomedical imaging data, leading to widespread use in clinical and research settings and a rich software ecosystem. By specifying a set of standards for storing metadata pertaining to image acquisition in file headers, and because DICOM is used nearly universally, it is possible to organize biomedical imaging data across sites using metadata tags. However, many DICOM tags pertaining to acquisition allow for the use of character strings, leading to arbitrary data entry that varies from site-to-site or between protocols. This complicates organizing MRI data using DICOM tags, despite the structure the standard provides.

While DICOM standardizes the storage of imaging data and metadata, there still exists a need to standardize analyses and derived results. In neuroimaging, the Brain Imaging Data Structure^5^ (BIDS) is a specification detailing how imaging datasets should be organized in a filesystem, facilitating sharing and reproducibility of MRI data, with some repositories requiring data to be in BIDS.^6^ While BIDS has seen increasing adoption^7,8^ it is still less mature than DICOM, and the hurdle of transforming DICOM data to BIDS is a major barrier to adoption. Furthermore, BIDS is intended only to be a filesystem specification, and providing an ontological representation of MRI data is beyond its scope.

Conversely, the Radiological Society of North America provides the RadLex ontology for harmonizing radiology terms and data.^9^ Widely used clinically, RadLex contains several terms governing MRI with some ties to DICOM. However, RadLex is focused on radiology, with an emphasis on describing radiological findings and procedures. Additionally, RadLex is clinically oriented and does not cover analysis of MRI data. In contrast, the National Institute of Health (NIH) provides a repository of Common Data Elements (CDEs) that includes several terms for MRI data and analysis. CDEs are comprised of questions and responses with designated data types, and are used by the NIH for standardizing data collection in research. While CDEs provide standardized data elements that can be used in conjunction with ontology, they do not explicitly define the ontological relationships required for a comprehensive representation of MRI.

To address these problems, we have developed the MRI Ontology (MRIO), building upon the work of Serra et al., to establish a “common language” for MRI acquisition and analysis^10^. For MRI, an ontology helps to standardize the naming and definition of entities related to acquisition, analysis, and dataset transformation. Specifically, we have developed definitions for entities related to acquisition such as relaxation times, as well as specific acquisition types such as T1 weighted (T1w) and T2 weighted (T2w). Although MRIO is generalizable to MRI as a whole, the focus of the discussion will be on its applications in neuroimaging. MRIO provides a formal framework for planning and organizing neuroimaging studies and analyses from acquisition to image processing, based on a standard set of classes and relationships, allowing for interoperability and harmonization of data and analyses across sites. MRIO was developed in accordance with Open Biomedical Ontology (OBO) Foundry principles^11,12^ and has contributed several terms to the Ontology for Biomedical Investigations (OBI).^13^

## Methods

### Development

MRIO was developed in the Web Ontology Language (OWL) 2^14^ using Protégé 5.5.0^15^ in accordance with OBO Foundry principles ^11^. Development was managed using the Ontology Development Kit^16^ for dependency management and interoperability. Logical consistency of axioms was assessed by using the HermiT OWL 2 reasoner.^17^ MRIO, along with all code and scripts, is publicly available on GitHub (https://github.com/Buffalo-Ontology-Group/MRI_Ontology) and is covered by the Creative Commons BY 4.0 license.

### Construction

MRIO builds upon OBI and the Information Artifact Ontology (IAO),^18^ operationalizing MRI in a standardized format used in other domains. Most terms in MRIO are children of the *information content entity* (IAO:0000030), *data set* (IAO:0000100), and *data transformation* (OBI:0200000) classes. Additionally, we have contributed axioms to the *magnetic resonance imaging assay* (OBI:0002985) term from OBI to better fit our paradigm by providing axioms for an input, *protocol* (OBI:0000272), *evaluant role* (OBI:0000067), and output (Figure 1). Thus, we have expanded on the existing term to represent the process of acquiring MRI, with the possibility to specify different types of MRI – e.g. T1w, T2w – depending on the *protocol* and the scanning parameters inherent to that *protocol*.

**Figure 1.**
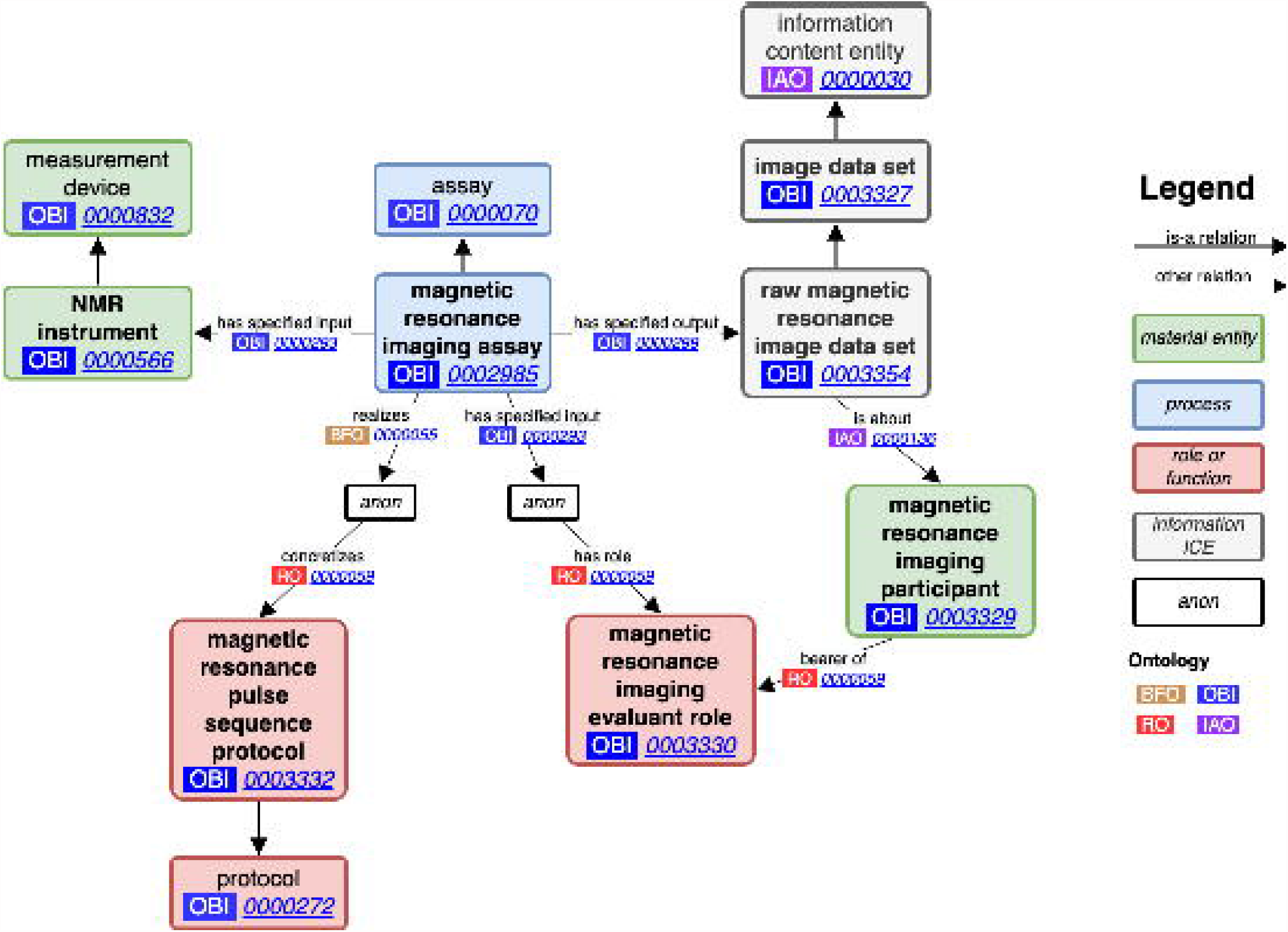
Magnetic resonance imaging assay: Our contributed terms and edits to the *magnetic resonance imaging assay* term from OBI (OBI:0002985). We added axioms for the pulse sequence protocol used in MRI acquisition, as well as an evaluant role used to tie back to the *raw magnetic resonance image data set* (OBI:0003354) output via its relation to the *magnetic resonance imaging participant* (OBI:0003329).

Because the physics of MRI acquisition is complex, there is no consensus on exact parameters defining what the acquisition of different MRI types should be. In reality, there are often ranges of values for parameters that may be encoded into a protocol on an MRI machine for individual MRI acquisition types. To define these ranges for each acquisition type, we combined domain experience and literature review with analysis of scans from public datasets and data from past studies.^19^ The public datasets analyzed were the Alzheimer Disease Neuroimaging Initiative (ADNI)^20^ and the Parkinson Progressive Marker Initiative (PPMI).^21^ This analysis used the pydicom ^22^ library to extract the range of values for the following commonly reported acquisition parameters: echo time (TE), repetition time (TR), inversion time, flip angle, field of view, slice gap, slice thickness, phase encoding direction, echo train length, and scanning sequence – e.g. spin echo, gradient echo, echo planar, and inversion recovery (Table 1).

**Table 1.** MRI acquisition parameters.

| **Class name** | **IRI** | **Definition** |
| --- | --- | --- |
| 'MRI machine parameter' | MRIO:0000342 | Parameters that a technician may specify to compose a specific MRI acquisition sequence as part of a scanning protocol. |
| 'acquisition matrix' | MRIO:0000364 | The total number of independent data samples in the frequency and phase directions (number of pixels in the frequency encoding - and phase encoding direction). |
| 'diffusion b value' | MRIO:0000655 | Measures the degree of diffusion weighting applied, thereby indicating the amplitude (G), time of applied gradients ($\delta$) and duration between the paired gradients ($\Delta$). |
| 'diffusion gradient' | MRIO:0000690 | Magnetic field variations applied to determine directional vectors that measure at least three dimensions of diffusion for a given voxel. |
| 'echo time' | MRIO:0000358 | The tie between an excitation pulse and peak of the echo produced in magnetic resonance imaging. |
| 'echo train length' | MRIO:0000359 | The number of spin echoes in given unit of time. |
| 'field of view' | MRIO:0000360 | The distance (in cm or mm) over which a magnetic resonance image is acquired or displayed. |
| 'flip angle' | MRIO:0000630 | Angle at which a proton's axis shifts from its longitudinal place to its transverse plane by excitation from radiofrequency pulses. |
| 'inversion time' | MRIO:0000653 | The time elapsed between the preparatory 180° pulse and the 90° readout pulse. |
| 'matrix size' | MRIO:0000369 | The number of data points collected in one, two or all three directions. Normally used for the 2D in plane sampling. |
| 'number of temporal positions' | MRIO:0000687 | Prescribed timepoints to collect in a 4D scan, results in in n number of volumes. |
| 'phase encoding' | MRIO:0000365 | The process of locating a magnetic resonance signal by altering the phase of spins in one dimension with a pulsed magnetic field gradient along that dimension prior to the acquisition of the signal. |
| 'phase FOV' | MRIO:0000361 | A parameter that can be selected by a technologist as a percent of our overall field of view. A different field of view (the scanned region) in the frequency and phase encoding directions that means the data acquisition with fewer measurement lines. |
| plane | MRIO:0000356 | The direction that the MRI image was taken. |
| coronal plane | MRIO:0000371 | The plane that divides the body into ventral and dorsal sections. |
| sagittal plane | MRIO:0000370 | The plane that divides the body into left and right sections. |
| transverse plane | MRIO:0000372 | The plane that divides the body into inferior and superior sections. |
| 'repetition time' | MRIO:0000357 | The repetition time (TR) is the time from the application of an excitation pulse to the application of the next pulse. It determines how much longitudinal magnetization recovers between each pulse. It is measured in milliseconds. |
| 'slice gap' | MRIO:0000363 | The not measured distance between the slices. |
| 'slice thickness' | MRIO:0000362 | The distance between two 2D image matrices, gives the measurement of a voxel in the z direction. |

Likewise, analysis aimed at deriving neuroinformatic data is complex and does not lend itself to be captured in a single ontology. Neuroimaging analysis may be thought of broadly as pipelines of coordinated software that are themselves often modular assemblies of imaging analysis algorithms. Many analysis pipelines are built on a combination of well-known and peer reviewed software suites such as the FMRIB Software Library (FSL), ^23^ Analysis of Functional NeuroImages (AFNI) ^24^, and Statistical Parametric Mapping (SPM).^25^ Rather than attempting to represent all combinations of pipelines created from these software, we instead represent the most common kinds of analysis pipelines agnostic of the software used. In some cases, however, terms were added for specific software as examples of analyses or where the software is ubiquitous with the analysis type – e.g. FreeSurfer^26–28^ for cortical thickness estimation. Table 2 contains a breakdown of the software included in MRIO.

**Table 2.** Neuroimaging software terms in MRIO.

| **Analysis** | **Definition** | **DOI** |
| --- | --- | --- |
| 'DeepGRAI analysis' | A volumetric magnetic resonance imaging analysis that uses a neural network (U-net) to estimate thalamic volume from T2 FLAIR image datasets of an entire human brain. | https://doi.org/10.1016/j.nicl.2021.102652 |
| 'FreeSurfer analysis' | FreeSurfer is a suite MRI data set analysis software that automates tissue segmentation and volumetry estimates from T1w MRI. | https://doi.org/10.1016/j.neuroimage.2012.01.021 |
| 'FreeSurfer cross-sectional analysis' | FreeSurfer tools to derive surface labels and volumes from an individual at only one timepoint. |  |
| 'FreeSurfer longitudinal analysis' | A set of FreeSurfer tools designed to automate the quantification of changes in brain morphometry over time from T1w MRI of an individual taken at several time points. |  |
| 'NeuroSTREAM analysis' | A brain volumetric analysis that automates estimation of lateral ventricle volume from clinical quality T2w FLAIR MRI. | https://doi.org/10.1016/j.nicl.2017.06.022 |
| 'SIENAX analysis' | Brain volumetric analysis tool for the automated estimation of grey matter, white matter, CSF, and whole brain volume from T1w MRI. | https://doi.org/10.1006/nimg.2002.1040 |
| 'SIENA analysis' | A brain volumetric analysis tool that estimates the percent brain volume change in an individual between two timepoints using baseline and follow-up T1w MRI. | Smith, Stephen M.; De Stefano, Nicola; Jenkinson, Mark; Matthews, Paul M. Normalized Accurate Measurement of Longitudinal Brain Change, Journal of Computer Assisted Tomography: May 2001 - Volume 25 - Issue 3 - p 466-475 |
| 'LPA analysis' | A focal lesion analysis tool that uses statistical methods to predict the location of MS lesions in a T2w MRI. | https://doi.org/10.5282/edoc.20373 |
| 'c-pac analysis' | A functional MRI data set analysis software suite of preconfigured pipelines for analysis fMRI data and estimating functional connectomics. | https://doi.org/10.3389/conf.fninf.2013.09.00042 |
| 'FAST analysis' | An MRI data set analysis tool used for segmenting and estimating the volume of grey matter, white matter, and CSF | Zhang, Y. and Brady, M. and Smith, S. Segmentation of brain magnetic resonance images through a hidden Markov random field |
|  | from T1w MRI. | model and the expectation-maximization algorithm. IEEE Trans Med Imag, 20(1):45-57, 2001. |
| 'FIRST analysis' | An MRI data set analysis tool for automated segmentation and volume estimation of subcortical structures (e.g. thalamus, cuneus, etc.). | Patenaude, B., Smith, S.M., Kennedy, D., and Jenkinson M. A Bayesian Model of Shape and Appearance for Subcortical Brain NeuroImage, 56(3):907-922, 2011. |
| 'NeMo analysis' | The NeMo Tool uses a large reference set of healthy tractograms to assess implied network changes arising from a particular pattern of WM alteration on a region- and network-wise level. | https://dx.doi.org/10.1089%2Fbrain.2013.0147 |

## Results

### MRIO classes and relations

Table 3 provides a breakdown of the number of axioms, classes, and object and data properties developed for MRIO. To date, 9 terms have been contributed to OBI and imported back to MRIO, with 28 terms in the process of being added to OBI. This section summarizes top-level classes in MRIO, using **bold face** to introduce each term and *italics* to denote a class, instance, or relation.

**Table 3.** Ontology metrics.

| **Metric** | **Count** |
| --- | --- |
| Axioms | 1,769 |
| Logical axioms | 511 |
| Declaration axioms | 323 |
| Classes | 279 |
| Object properties | 18 |
| Data properties | 6 |

#### Image data set

MRI data is a special kind of digital “image” that typically represents three-dimensional space using arrays of voxels, similar pixels in two-dimensional images. Some MRI even use four-dimensional arrays, like functional or diffusion MRI. Additionally, biomedical imaging data contain metadata pertaining to its acquisition in DICOM headers, which are vital for understanding the type of data.

As such, we needed a generalizable term for MRI data, as one did not already exist in OBO Foundry ontologies. There is *image* (IAO:0000101), but it has a rigid definition as two-dimensional and does not capture the data in DICOM headers pertaining to acquisition. To provide a better representation of biomedical imaging data, we developed the *image data set* class (OBI:0003327) as a subclass of *data set* (IAO:0000100) with the following definition: “A data set that is comprised of multidimensional structured measurements and metadata required for a morphological representation of an entity.”

#### Image data set analysis

In order to fully capture the neuroimaging research process, we needed a generalizable term representing the analysis of MRI data. Neuroimaging analysis is a broad field that involves transforming entire *image data sets* into new *image data sets* or deriving numeric metrics. We needed to capture the range of data that may be collected from an *image data set*, with a broad definition that allows for the different kinds of output created.

Therefore, we developed *image data set analysis* (OBI:0003355) as a child of *data transformation* (OBI:0200000) with the following definition: “The process of deriving a data item from an image data set using computer algorithms.” The term *image data set analysis* is axiomatically defined as the subclass of *data transformation* that *has_specified_input* some *image data set*.

#### Raw image data set

We developed two subclasses for *image data set* to represent the raw data acquired by an imaging device and the transformed data that is more commonly used in analyses. For the former case we developed *raw image data set* (OBI:0003327) to represent the kinds of raw data acquired by an MRI scanner before it is transformed into the more common DICOM or NIfTI formats. We have defined *raw image data set* as follows: “An image data set that encodes measurement values produced by some instrument before undergoing a data transformation.”

#### Raw magnetic resonance image data set

An MRI scanner uses a series of magnetic gradients and radiofrequency pulses to enable spatial resolution of the tissue being examined based on the frequencies and phases of the signals the scanner receives from the tissue. This raw signal data is organized in k-space, a special kind of *image data set* that is unsuitable for analysis or clinical use, but is nevertheless an integral part of MRI acquisition. We developed *raw magnetic resonance image data set* (OBI:0003354) as a subclass or *raw image data set* to capture data with the following definition: “An image data set that is the direct output of a magnetic resonance imaging assay and whose values encode spatial frequencies produced by the NMR or MRI instrument.”

#### Computed image data set

Most imaging data that researchers and interact with is the product of some mathematical transformation applied to the raw data produced by the imaging hardware to better visualize the morphological representation of an entity. In the case of MRI, a Fourier transform is often applied to the raw data to produce the three-dimensional volume in the shape of the tissue being examined. We developed *computed image data set* (OBI:0003333) to represent these data with the following definition: “An image data set that is the output of an image data set analysis.” A *computed image data set* is axiomatically defined as the *specified output* of some *image data set analysis*.

#### Reconstructed magnetic resonance image data set

Most MRI data is stored and analyzed as DICOMs or NIfTIs consisting of volumes of voxel arrays that visually represent some tissue in three-dimensional space. This kind of *image data set* is the result of a mathematical transformation of the raw k-space data into a multi-dimensional array called a “reconstruction.” To represent these data, we developed *reconstructed magnetic resonance image data set* (OBI:0003334) as a subclass of *computed image data set* with the following definition: “An image data set that is the direct output of a raw magnetic resonance image data set reconstruction or a transformation of another magnetic resonance image data set.”

Because *reconstructed magnetic resonance image data set* represents most MRI data, we developed subclasses of *reconstructed magnetic resonance image data set* corresponding to common MRI types. These include *T1 weighted magnetic resonance image data set* (MRIO:0000366) and *T2 weighted magnetic resonance image data set* (MRIO:0000367), as well as child classes for specific types such as *T1 SPGR magnetic resonance image data set* (MRIO:0000680) for the high-resolution MRI used in academic research and *T2 FLAIR magnetic resonance image data set* (MRIO:0000511) for the modality used in the clinical routine of MS. Using these subclasses, we organized common MRI types into well-reasoned groups according to their acquisition parameters (Figure 2).

**Figure 2.**
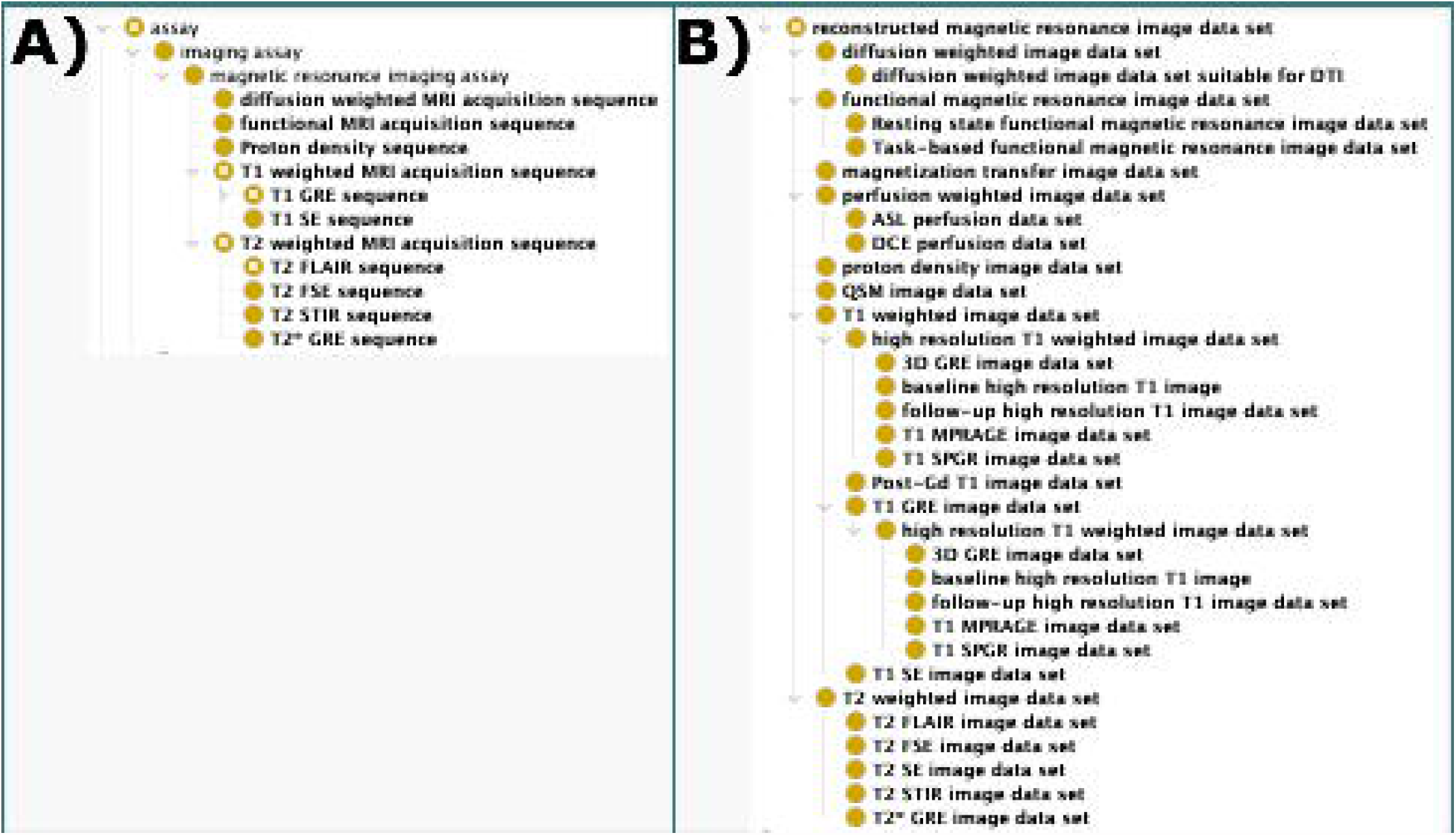
MRI acquisition types: The acquisition of MRI data from a scanner is represented in MRIO in two steps: the imaging assay that describes the process and the image data set that describes the data itself. A) Different MRI acquisition types are defined according to their acquisition parameters, each with the own logical and semantic definitions. We group similar MRI acquisition types together (i.e. T1 weighted MRI is commonly used to refer to T1 weighted images acquired using spin echo (SE) or gradient echo (GRE) sequences). B) Reconstructed MRI are often the output of specific MR imaging assay sequences, while some are produced by other MRI analyses and data transformations. These classes represent the data that is used clinically or analyzed in the research setting.

MRI types are defined by the acquisition sequence used at the scanner. At a high level, MRI are acquired using a magnetic field to align protons in the examined object, a radiofrequency pulse to move protons from alignment, and a receiving coil to pick up signals given off by protons as they move back through the magnetic field into alignment. MRI types are divided into groups depending on the time it takes protons to return to alignment (i.e. “relaxation time”). Relaxation time largely controls contrast, which is a factor of the T1 “spin-lattice” and T2 “spin-spin” relaxation times. These form the two largest groups, T1w and T2w MRI, which show differing contrast in the same tissue and are acquired by tuning the scanner to interrogate properties of the underlying tissue, including the TE and TR. TE is a measure of the time between the application of the radiofrequency pulse and when the peak signal is received, while TR is the time between radiofrequency pulses. We created a data property to represent the numerical properties used in an MRI acquisition sequence called *has MRI machine setting value* with the following definition: “A relation between the MRI machine setting datum set by the technician and the properties of an MRI acquisition sequence.”

T1w MRI are acquired using lower TE and TR, while T2w MRI use higher TE and TR. We created two classes that represent T1w and T2w MRI based on their acquisition parameters, then subdivide them based on additional acquisition parameters. We define a *T1 weighted MRI acquisition sequence* (MRIO:0000385) having a *T1 weighted image data set* (MRIO:0000366) as its specified output with the following logical axioms: (*has TR value* some xsd:float[> 0.0f, < 800.0f]) and (*has TE value* some xsd:float[> 0.0f, < 30.0f]) and (*has flip angle value* = 90). Conversely, we define a *T2 weighted MRI acquisition sequence* (MRIO:0000386) having a specified output of a *T2 weighted image data set* (MRIO:0000367), with the following logical axioms: (*has TR value* some xsd:float[> 2000.0f]) and (*has TE value* some xsd:float[> 30.0f]) and (*has flip angle value* = 90). We also developed terms for the acquisition of DWI, fMRI, and proton density (PD) MRI as subclasses of *reconstructed magnetic resonance image data set*.

#### Magnetic resonance image data set analysis

Neuroimaging analysis is a broad field, answering questions about the brain from multiple disciplines and using different MRI types. To address this, we developed a general class as a child of *image data set analysis* called *magnetic resonance image data set analysis* (MRIO:0000508) with the following definition: “Any combination of digital image analysis and statistical tools that may be used to derive processed magnetic resonance images or data measurements about the magnetic resonance images,” with a logical axiom of (*has_specified_input* some *reconstructed magnetic resonance image data set*). There is no specified output for *magnetic resonance image data set analysis* as it may be used to derive any metric from the MRI data set or produce a transformed version of the MRI data set. Additionally, several subclasses of *magnetic resonance image data set analysis* have been developed to represent the kinds of analyses that may be done and the MRI type that an analysis requires as input (Figure 3). These analyses are grouped according to the information they derive from the brain or transformation the MRI undergoes, with several software added as their own subclasses (Table 2, Figure 4).

**Figure 3.**
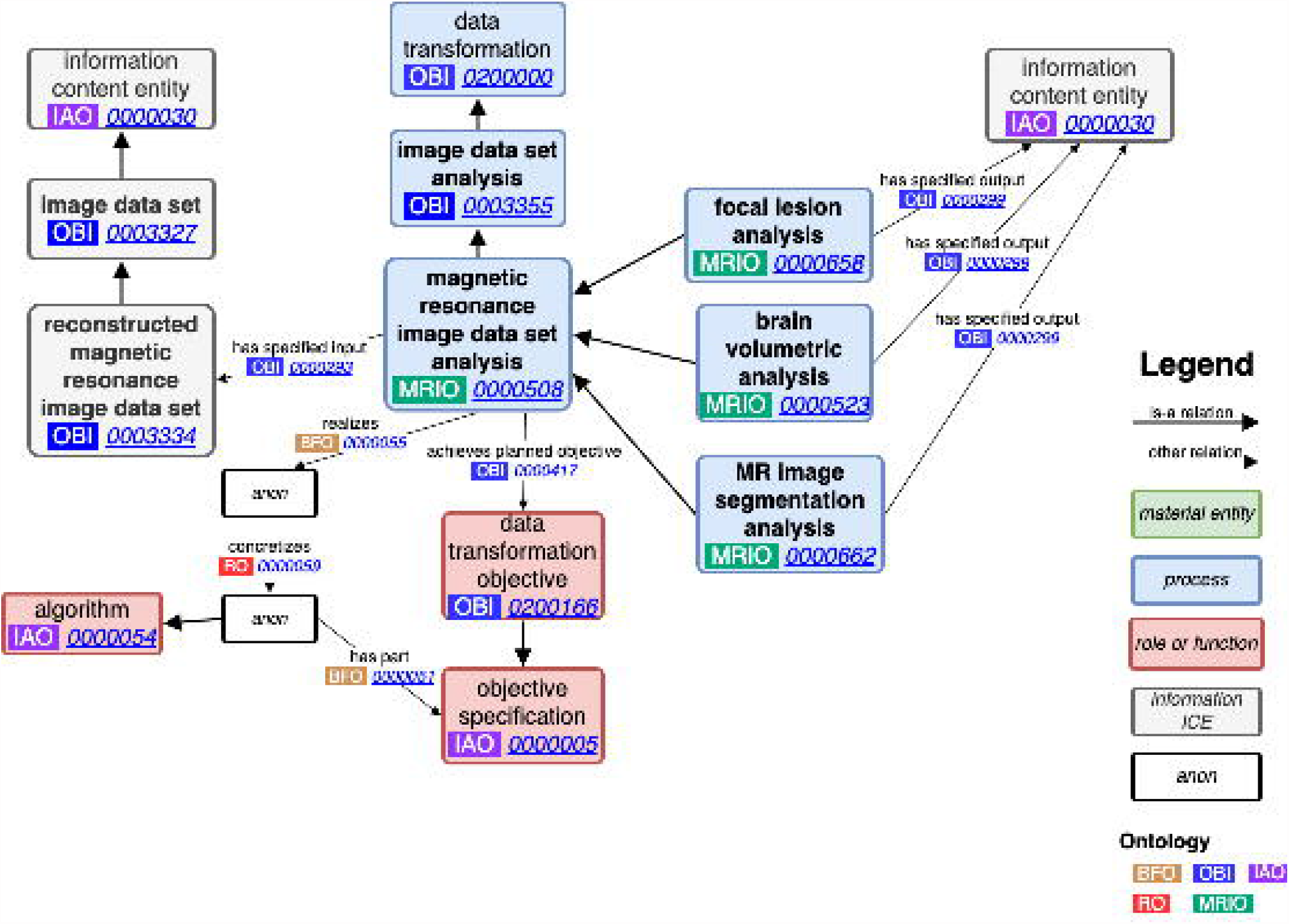
Magnetic resonance image data set analysis: The process of analyzing MRI data follows the format of OBI’s data transformation, applying algorithms to reconstructed magnetic resonance image data sets to produce some information content entity. The output could be a specific value or even a new, transformed magnetic resonance image data set.

**Figure 4.**
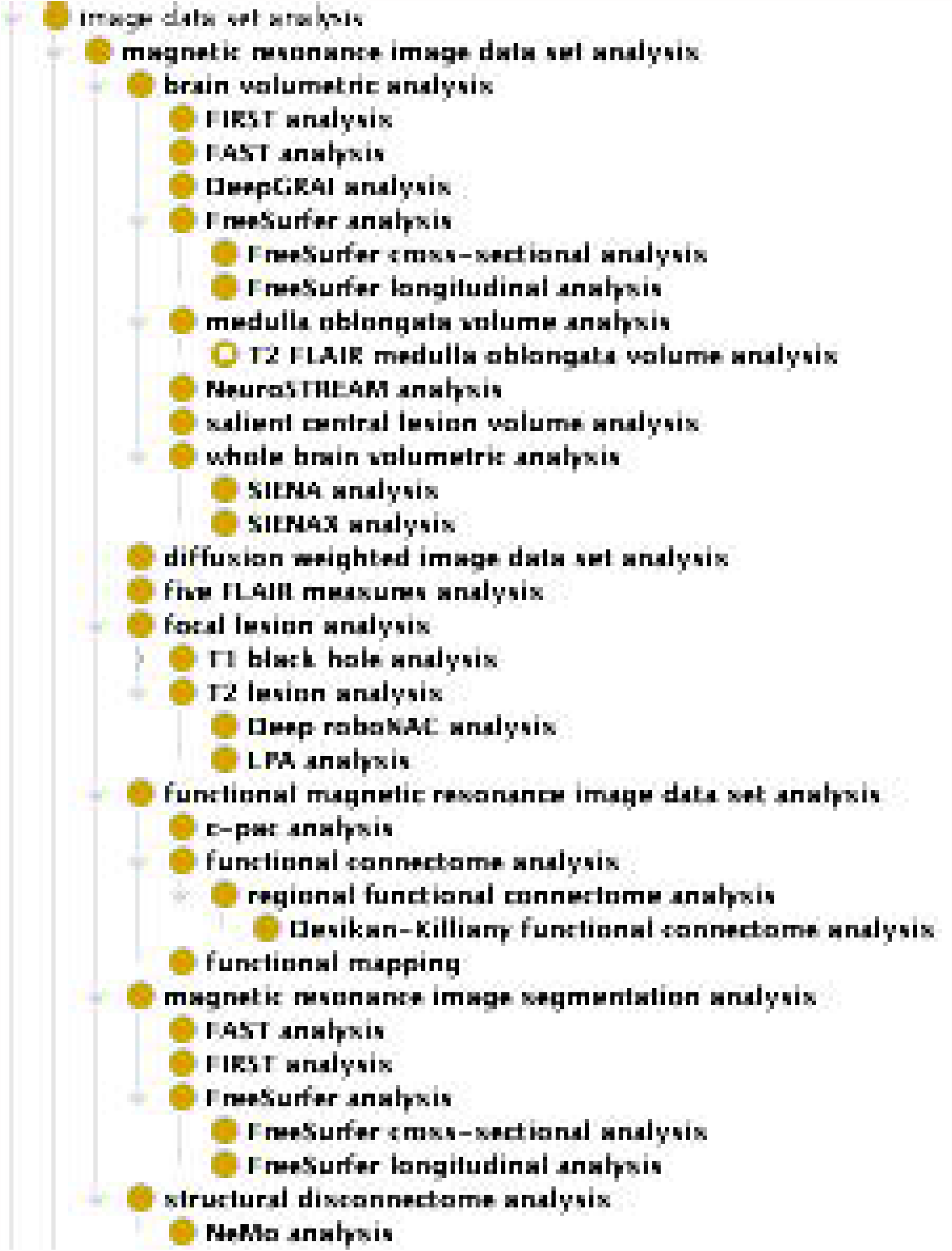
MRI analysis terms: MRIO contains groups of common neuroimaging analyses types, with specific analyses included as child classes. Specific MRI analyses are grouped by similar processes or the regions of the brain they analyze.

For example, *SIENAX analysis* (MRIO:0000525) is a subclass of *whole brain volumetric analysis* (MRIO:0000524), which is in turn a subclass of *brain volumetric analysis* (MRIO:0000523), which is one of the analysis groupings under *magnetic resonance image data set analysis. SIENAX analysis* has the following logical axioms (*has_specified_input* some (*T1 weighted image data set* and *T2 FLAIR image data set*) or *has_specified_input* some *T1 weighted image data set*), (*has_specified_output* some (*whole brain volume measurement datum* and *grey matter volume measurement datum* and *white matter volume measurement datum* and *CSF volume measurement datum*)). Using these axioms, we are able to fully represent the kind of data required to run FSL’s SIENAX software and the metrics that are derived from it.

#### Brain region atlas image data set

In MRI analysis, it is necessary to use standardized templates or atlases that encode the coordinates of specific brain regions. We developed a class for these called *brain region atlas image data set* (MRIO:0000647), which is “An image data set consisting of values computed from multiple image data sets encoded to represent the spatial location of individual functional or structural regions of a canonical brain,” with the intent that specific brain atlases be added as children. For example, we also developed the *Desikan-Killiany brain atlas image data set* (MRIO:0000649) for FreeSurfer and added *is about* relations to each of its 86 regions atlas, mapping them to their corresponding IRIs in Uberon.^29^ This helps to standardize reporting on analyses using these brain regions and tie neuroinformatics into existing databases that use Uberon.

## Discussion

Here we report on the development of MRIO, the MRI ontology built according to OBO Foundry principles. Our work addresses the need for standardized and interoperable representation of MRI data, providing a framework for capturing the intricacies of acquisition and analysis. With several terms contributed to OBI, MRIO facilitates integrating MRI data across biomedical domains. By formalizing the concepts and relationships in MRI analysis, MRIO offers a valuable resource for researchers and clinicians working with MRI data. MRIO makes managing the neuroimaging research process easier by helping automate the organization of MRI data, automatically assigning analyses based on available MRI acquisition types, and integrating with existing standards.

One of the hurdles researchers encounter working with MRI data is determining what MRI types are in their dataset, what analyses can be done, and what the results mean. By creating terms for common MRI types and analyses and organizing them into standardized, we developed a simple tool to automatically assign common analyses to a given dataset depending on the MRI types available, which is available in the MRIO repository. The tool takes an array of MRI type IRIs and chains several SPARQL Protocol and RDF Query Language (SPARQL) queries together that find every possible analysis that takes that MRI type as an input. The first query filters matching MRI types and uses those as the input for the next query. The second query identifies analysis terms that contain an axiom for accepting only a single available MRI acquisition type as an input, while a third query matches analysis terms that accept multiple available MRI acquisition types. The tool returns a dictionary of MRIO IRIs mapped to their labels for all available analyses that may be performed on a given list of MRI types.

The structure of classes representing MRI types and analyses in MRIO are intentionally similar to the filesystem and naming conventions used by BIDS. Furthermore, the acquisition parameters used in the logical axioms of MRI types in MRIO match the parameters reported in an MRI’s sidecar JSON file used by BIDS (e.g. TE, TR, etc.). This allows for integration with the BIDS ecosystem, helping researchers use standardized formats with their imaging data. More specifically, MRIO provides a means for harmonizing the annotation of both DICOM and BIDS datasets by providing a unique identifier that maps corresponding DICOM header tags to JSON keys in BIDS sidecars. For example, the value for TE used to acquire an MRI is encoded in DICOM headers with Tag (0018,0081), while BIDS will use “EchoTime” as a string formatted key in its sidecar. MRIO provides the IRI MRIO:0000358 for *echo time*, bridging the gap between DICOM and BIDS representations of acquisition and ensuring consistent and standardized annotation (Figure 5). This aids researchers in transforming raw data into organized BIDS datasets. There is also potential for MRIO to enrich existing software used for generating BIDS datasets from DICOM, such as BIDScoin^30^ or CuBIDS, ^31^ which require that a user supplies ranges for acquisition parameters as heuristics to sort available MRI types into BIDS. Since MRIO supplies logical axioms containing ranges for MRI acquisition parameters, queries of MRIO can help automate the transformation of unsorted DICOMs into BIDS.

**Figure 5.**
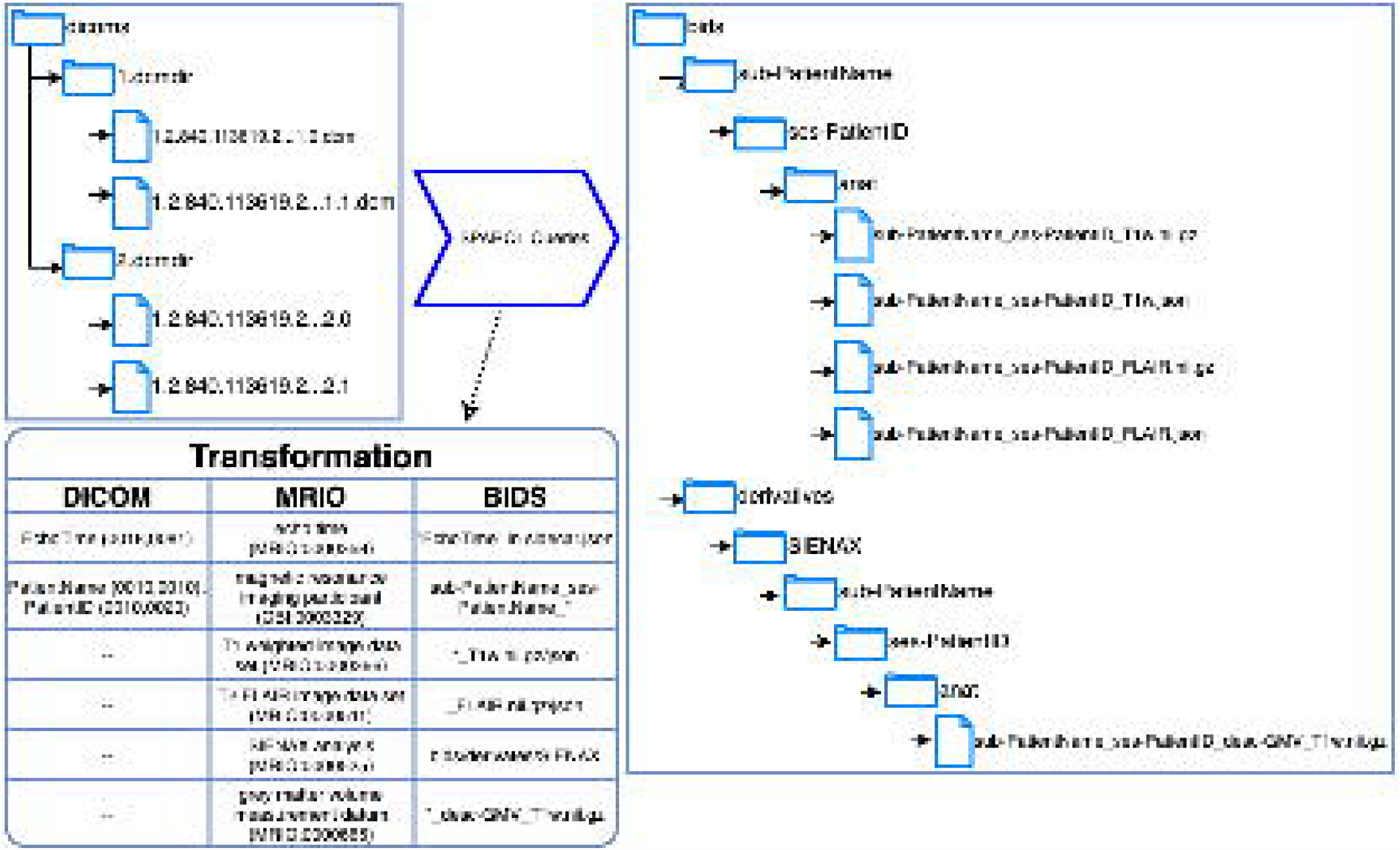
DICOM to BIDS transformation: BIDS specifies directory structures and naming conventions for storing MRI data on a file system, making BIDS datasets easier for a human to read and work with than raw DICOMs. MRIO can serve as a middle layer between DICOM and BIDS datasets, with terms that map standardized DICOM tags to the naming conventions specified by BIDS. Additionally, analysis terms from MRIO – including the name of the analysis and kind of data produced – can be used to annotate analyses of BIDS data sets, which BIDS call “derivates.”

One of MRIO’s key functions is the annotation of MRI databases with essential information such as acquisition parameters, types, analyses, and results. This process enhances the organization and discoverability of data, helping researchers to retrieve subsets based on their characteristics. By using reasoners leveraging the knowledge captured in MRIO, researchers can streamline the analysis workflow by programmatically assigning analyses. This reduces the effort involved in managing and processing large datasets, saving time and enhancing efficiency, enabling researchers to extract meaningful insights from vast amounts of MRI data.

MRIO was developed in accordance with OBO Foundry principles and has contributed several terms to OBI. As such, MRIO integrates with a wide range of ontologies across biomedical domains. This integration is already present in MRIO, with axioms used to define brain atlases that map to anatomical regions defined in Uberon. There is also potential to harmonize MRI data across adjacent fields with preestablished ontologies, such as clinical research using the Neurological Disease ontology^32^ and Neurodegenerative Disease Data Ontology.^33^ By building on and contributing to OBI, there is also potential to integrate with existing MRI ontologies, such as RadLex or the Neuroimaging Data Model, ^34,35^ enhancing the impact of both and reducing duplication of effort.

While MRIO makes strides in harmonizing MRI, we recognize there remains the need for a fuller representation of this domain. First, neuroimaging is a broad field that is constantly evolving with many more MRI types and analyses than are captured in MRIO. Our work focuses on MRI types common to the clinical routine. As such, there are undoubtably several MRI types and analyses missing from MRIO that would be useful for many in the community, such as field mapping, susceptibility weighted imaging, etc. Our intent was to provide a strong core of commonly encountered MRI types and analyses, with the hope that the open-source nature of MRIO enables others in the community to contribute terms as the field evolves. Secondly, while the axioms used to define MRI acquisition sequences are based on many years of domain experience and thorough analysis of large public MRI datasets, there is an inherent limit to the generalizability of the range of values provided by MRIO. There are no hard-and-fast rules for how MRI types are acquired – the contrast is a function of the MRI scanner, parameters used, and properties of the tissue examined. Nevertheless, we provide what we believe are reasonable ranges for common acquisition parameters. Future work should incorporate more datasets to round out the ranges used to define acquisition in MRIO. Finally, there is need to incorporate feedback from the broader neuroimaging community, extending the impact of MRIO.

## Conclusion

MRIO provides researchers a means of managing the neuroimaging research process. By developing a framework for annotating and organizing MRI data, MRIO helps promote the reproducibility and interoperability of neuroimaging studies. By providing well-reasoned logical definitions for common MRI types and analyses, MRIO reduces the barrier to entry for clinical and translational researchers hoping to work with MRI data. Built according to OBO Foundry principles, MRIO integrates with a diverse community of biomedical ontologies, helping bridge the gap between neuroimaging and other domains.

## Acknowledgements

The MRIO team would like to thank the developers of OBI – in particular James Overton, Bjoern Peters, and Chris Stoeckert – for their contributions and guidance as we prepared our higher-level terms for the broader biomedical ontology community. We are grateful for the opportunity to contribute our work to such a foundational ontology.

